# ELIMINATOR: Essentiality anaLysIs using MultIsystem Networks And inTeger prOgRamming

**DOI:** 10.1101/2021.07.21.453265

**Authors:** Asier Antoranz, María Ortiz, Jon Pey

## Abstract

A gene is considered as essential when it is indispensable for cells to grow and replicate under a certain environment. However, gene essentiality is not a structural property but rather a contextual one, which depends on the specific biological conditions affecting the cell. This circumstantial essentiality of genes is what brings the attention of scientist since we can identify genes essential for cancer cells but not essential for healthy cells. This same contextuality makes their identification extremely challenging. Huge experimental efforts such as Project Achilles where the essentiality of thousands of genes is measured in over one thousand cell lines together with a plethora of molecular data (transcriptomics, copy number, mutations, etc.) can shed light on the causality behind the essentiality of a gene in a given environment by associating the measured essentiality to molecular features of the cell line.

Here, we present an in-silico method for the identification of patient-specific essential genes using constraint-based modelling (CBM). Our method expands the ideas behind traditional CBM to accommodate multisystem networks, that is a biological network that focuses on complex interactions within several biological systems. In essence, it first calculates the minimum number of non-expressed genes required to be active by the cell to sustain life as defined by a set of requirements; and second, it performs an exhaustive in-silico gene knockout to find those that lead to the need of activating extra non-expressed genes.

We validated the proposed methodology using a set of 452 cancer cell lines derived from the Cancer Cell Line Encyclopedia where an exhaustive experimental large-scale gene knockout study using CRISPR (Achilles Project) evaluates the impact of each removal. We also show that the integration of different essentiality predictions per gene, what we called Essentiality Congruity Score, (derived from multiple pathways) reduces the number of false positives. Finally, we explored the gene essentiality predictions for a breast cancer patient dataset, and our results showed high concordance with previous publications.

These findings suggest that identifying genes whose activity are fundamental to sustain cellular life in a patient-specific manner is feasible using in-silico methods. The patient-level gene essentiality predictions can pave the way for precision medicine by identifying potential drug targets whose deletion can induce death in tumour cells.

## Introduction

We can define an essential gene as a gene whose activity is fundamental to sustain life (Bartha *et al*., 2018). It is precisely the critical importance of these genes that brings the attention of scientists. For instance, in cancer research, specific essential genes of this condition are considered as promising drug targets as their deletion can induce death in tumour cells (Tsherniak *et al*., 2017).

The essentiality of a gene is not a structural property, it depends on the biological scenario under consideration (Zhang and Lin 2009), including the cellular environmental conditions, disease phenotypes, etc. The context dependency of essential genes makes their experimental identification an extremely difficult task. The huge effort of experimental initiatives such as Project Achilles in creating an archive of essential genes is of utmost interest to the scientific community (Tsherniak *et al*., 2017). However, the biological context in which a particular gene turns out to be essential is exceptionally critical in cancer, where the essentiality of a gene could emerge at patient level (Fernald *et al*., 2011). This highlights the role of *in-silico* gene essentiality identification approaches that effectively integrate-omics datasets to contextualize a given biological scenario.

During the last decade, many successful examples have been presented on integrating omics datasets with biological networks in the context of efficient mathematical models to address an assortment of biomedical problems (Yan *et al*., 2018), including the identification of essential genes (Li *et al*., 2020). We can find relevant insights provided by these algorithms in different fields, ranging from microbiology (Plata *et al*., 2010) to cancer research (Frezza *et al*., 2011), among others.

Despite the recent advent of machine-learning based gene essentiality analyses (Kuang *et al*., 2020), (Schapke *et al*., 2020), traditionally, approaches referred to as Constraint-Based Modelling (CBM) leaded the field setting the foundations for the development of different methodologies to predict essential genes (Apaolaza *et al*., 2017), (Pey *et al*., 2017), (Tobalina *et al*., 2016). In essence, CBM integrates omics data in the context of genome-scale metabolic networks resulting in a linear system of inequalities. The arising system of inequations is usually solved using linear optimization techniques (Martin 2012). Here, essential genes emerge from their indispensability when ensuring the activity of an artificial metabolic reaction, referred to as biomass reaction, which involves the metabolic requirements of the cell for its replication (Agren *et al*., 2014).

In this work, we extend the ideas in traditional CBM by going beyond metabolism considering multisystem networks (Schaefer et al., 2009). In addition, and in analogy with CBM, here we identify genes whose activity is essential for a relevant biological task. Thus, the emerging set of essential genes will be richer and more diverse than in traditional CBM, capturing a variety of biological processes (Vaske *et al*., 2010).

Overall, this article introduces a new methodology for the *in-silico* identification of essential genes. This approach combines three main inputs: (i) An indispensable biological entity/process required to sustain cellular life, (ii) a set of interaction networks including the molecular requirements to activate the aforementioned indispensable entity and (iii) an experimental dataset that reflects, at least qualitatively, the genetic landscape of the sample/patient, *e.g*. gene expression data.

These inputs are subsequently encoded into a mathematical model (Integer Linear Program, ILP) (Schrijver, 1998) that finds the minimum number of non-expressed genes required to activate the given relevant function. Then, a systematic approach identifies artificial gene knockouts that lead to require additional unexpressed genes to activate the critical biological entity/process. These knockouts are precisely considered as **essential genes**. This is further illustrated in the manuscript through a series of toy examples. In addition, we successfully validated a continuous score representing the degree of essentiality of a given gene, referred to as the *Essentiality Congruity Score*. We also show the relevance of each of these inputs by evaluating the performance of the method in a different set of scenarios. Finally, we apply the methodology to a group of Breast cancer patients and subsequently support the relevance of the emerging essential genes based on a literature review.

## Methods

In the following section, we introduce the *in-silico* gene-essentiality framework presented in this article. In the first subsection, referred to as **Pathways**, we describe the biological pathway compendium used for the model; in the second subsection, mentioned as **Datasets**, we describe all the experimental data used throughout the study; the third subsection, **Mathematical Model**, describes the mathematical equations modelling the pathways and integrating the experimental data; and, in the fourth subsection, called **Gene Essentiality Analysis**, we present the pipeline that systematically find essential genes. Moreover, we present the *Essentiality Congruity Score*, which assigns a quantitative value to an otherwise binary score to represent the essentiality of a gene.

### Pathways

As in (Vaske *et al*., 2010), we consider a set of well-curated pathways from the (NCI-PID) (Schaefer et al., 2009) database, which are represented in the UCSC Pathway Tab Format. Vaske and co-workers provided further details about the characteristics of these pathways, including their consistency when capturing cancer related knowledge. In essence, these pathways comprise vertices and edges representing various types of biological entities and their interactions respectively. For instance, vertices could denote a gene/protein, gene complex or biological abstracts like “mitosis” or “cell motility”, among others, whilst edges represent activations/inhibitions or member/component associations (Vaske *et al*., 2010).

Following the UCSC Pathway Tab Format (Vaske *et al*., 2010), we will consider the following interactions: member (*member*>), component (*components*>), activation (-*a*>, -*t*>, -*ap*>) and inhibition (-*t*|,-*ap*|, -*a*|). As will be introduced in the next subsection, each one of these interactions is modelled by a specific set of mathematical equations.

### Datasets

We initially show the behaviour of the algorithm in a toy-example simulated dataset using the *Wnt receptor signaling pathway, planar cell polarity pathway*, from the NCI-PID which is shown in **Figure 3** (Schaefer *et al*., 2009).

Secondly, we applied the methodology in the set of pathways from the NCI-PID (Schaefer et al., 2009) using the gene expression data from the Cancer Cell Line Encyclopedia (CCLE) (Barretina et al., 2012) and validated the biological relevance of the predictions using the essentiality scores from the Achilles project (Dempster et al., 2019). Project Achilles is a systematic effort aimed at identifying and cataloguing gene essentiality across hundreds of genomically characterized cancer cell lines. These gene essentiality scores are obtained from CRISPR knockouts (CERES method) (Meyers et al., 2017) on several of the cells lines included in the Cancer Cell Line Encyclopedia (CCLE) (Barretina et al., 2012), a compilation of gene expression, chromosomal copy number and massively parallel sequencing data from nearly 1,000 human cancer cell lines. The gene expression data for the cell lines was obtained from the Gene Expression Omnibus (GSE36133) which includes 917 cell-lines annotated with 23521 gene identifiers (HGNC format). Gene expression data was binarized (1 expressed, 0 not expressed) using The Gene Expression Barcode 3.0 (McCall et al., 2012), (McCall et al., 2014). Probes were mapped to HGNC identifiers (GPL570, Affymetrix Human Genome U133 Plus 2.0 Array). The Achilles Essentiality Scores were downloaded from the DepMap portal (https://depmap.org/portal/download/, version 20Q1) which contained essentiality scores for 18333 genes in 739 cell lines, 478 of which were in common with the CCLE. In total, 1660 HGNC IDs were in common between Achilles, CCLE and the genes/proteins present in the NCI-PID pathways. The Achilles database included in DepMap contains missing values for several of these genes (NA values). After removing NA values, 26 cell lines didn’t contain any data for the 1660 genes in common, thus reducing the number of included cell lines to 452. Achilles scores represent gene essentiality, the more negative the score, the more essential the knockout of the gene is for a given cell-line.

Finally, we applied the gene essentiality method to Breast Cancer patient samples (Maubant S, 2012), (Maire V, 2013). This dataset includes transcriptome analysis of 130 breast cancer samples (41 TNBC; 30 Her2; 30 Luminal B and 29 Luminal A), 11 normal breast tissue samples and 14 TNBC cell lines. This dataset contains 178 array samples. 153 arrays were used to analyse 130 unique breast cancer samples from as many patients and 23 technical duplicates. In addition, 11 “Normal” samples from healthy breast tissue obtained from mammoplasty are included, as well as a collection of 14 breast cancer cell lines. Data production involved different array batches and hybridation series which were accounted for in the pre-processing of the data. Processed gene expression data and sample meta-data was obtained from the Gene Expression Omnibus (GSE65194). Samples belonging to cell lines were removed from further analysis. Gene expression data was discretized using The Gene Expression Barcode 3.0 (McCall et al, 2012), (McCall et al, 2014). Probes were mapped to HGNC identifiers (GPL570, Affymetrix Human Genome U133 Plus 2.0 Array).

### Mathematical model

The network format described above can be translated into a series of Boolean rules. However, the inherent complexity of these rules grows exponentially when regular-sized pathways are considered. In this subsection, we present the Integer Linear Programming framework (ILP) able to contend with complex networks and capturing all the essence of the Boolean rules. The ILP constitutes the core of the methodology. In essence, it calculates the minimum number of non-expressed genes required to activate a given biological function necessary to sustain cellular life. The mathematical equations in the ILP arise precisely from the structure of the pathway and its different interactions (**Figure 1**).

**Figure 1 -.**
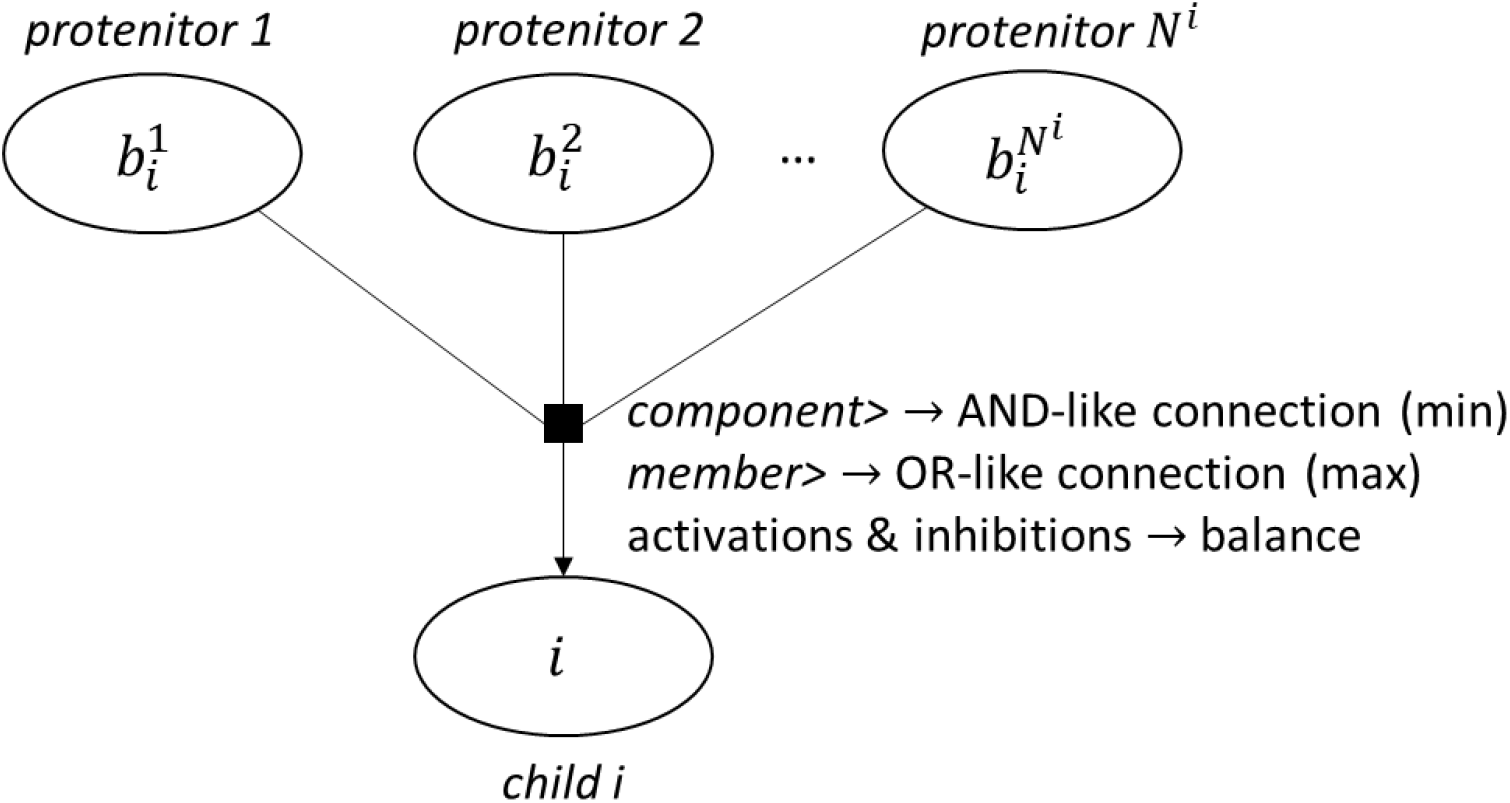
Conversion of UCSC Pathway Tab Format to valid pathways for the mathematical model. These pathways model the relationship between the i-th child and its progenitors using various types of interactions including component>, member>, and activations & inhibitions.

Let ***B^i^*** represent the set of all the parents for a given entity *i*. Let *N^i^* be the cardinality of ***B^i^***, namely *N^i^* = |***B^i^***|. We can define *E_i_* as a binary variable *{0,1}* that represents the final binary activation status of *i* in a given simulation (*E_i_* = 0 if inactive, *E_i_* = 1 if active). The reader should note that *Ei* is not the same as the experimental expression of the genes, the experimental data will be included later in the model. In particular, the method focuses on minimizing the number of non-expressed genes being active but does not apply direct restrictions to expressed genes.

Now we will proceed to mathematically define the constraints based on the nature of the interaction between the *i*-th child and its progenitors.

#### component>

In analogy with the AND-like connection considered for the components of a complex in Vaske *et al*., 2010, the final activation status of the child (*E_i_*) is determined by the minimum value from all its components. So *E_i_* = 1 if, and only if, *E_j_* = 1 *∀b* ∈ ***B_i_***. Otherwise, we impose that *E_i_* = 0.

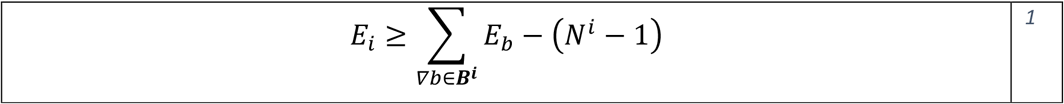

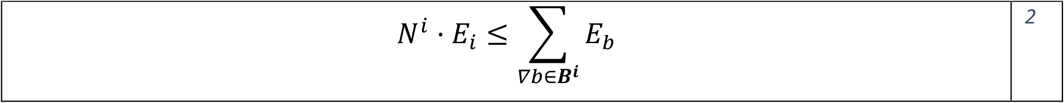

#### member>

The final status of node *i* is determined by the maximum value from all its members. So *E_i_* = 0 if, and only if, *E_b_* = 0 *∀b* ∈ ***B_i_***. Otherwise, we impose that *E_i_* = 1. Note the similarity with (Vaske *et al*., 2010) where members are modelled in an OR-like fashion.

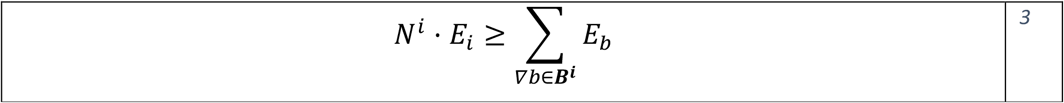

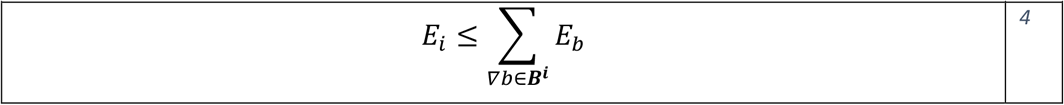

#### Activations & Inhibitions

The final status of the target is determined by the balance between all its activators and inhibitors. For simplicity, we can define an intermediate variable 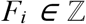 that expresses the activation/inhibition state of *i*,

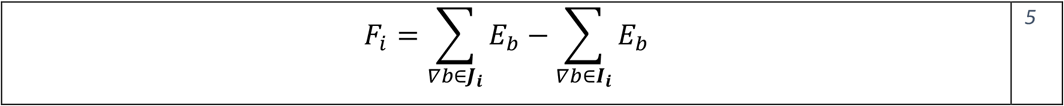

where ***J_i_*** and ***I_i_*** represent the set of activators and inhibitors progenitors of *i*, respectively. The activation status of the child *i* is then determined by its activation state,

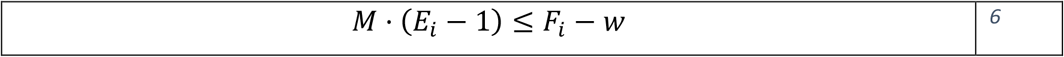

where *M* is an auxiliary positive large integer (*M* = 1000) and *w* the relative weight between activators and inhibitors that modules the sign of *F_i_*. Here, we considered an arbitrary value of *w* > 0,5. The reader should note how eq 6 forces *E_i_* = 0 when *F_i_* < *w* and does not constrain *E_i_* when *F_i_* ≥ *w*. That is, an inhibitory state of *i* is sufficient for the inactivation of the child, while an activation state of *i* is necessary for the activation of the child. Note that eq 6 is only imposed when the target *i* is an abstract or a complex because the genes and proteins generally represent the entries of the networks and their global activators-inhibitors scenario are often not properly captured in individual pathways.

#### Artificially activating an abstract/complex

We will impose the activation of relevant biological functions. To that end, we define the set of all entities required to sustain cellular life as ***A***, from now on defined as actives, and an independent problem is defined for each of them.

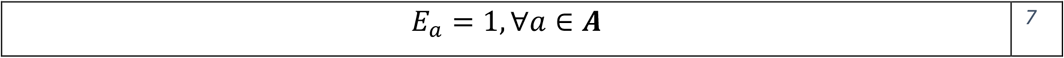

Where *E_a_* represents the activity of the entity *a*. In practice, for a given pathway, the set **A** consists of all its abstracts and complexes.

#### Minimizing the number of lowly expressed

Let ***L*** represent the set of non-expressed genes. ∀*a* ∈ ***A***, we define the optimal solution as the one that directly minimizes the number of non-expressed genes active in the final solution whilst *E_a_* = 1. Note that the model will provide a specific value of the objective function for each *a* ∈ ***A***. We will refer to this solution as 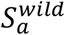.

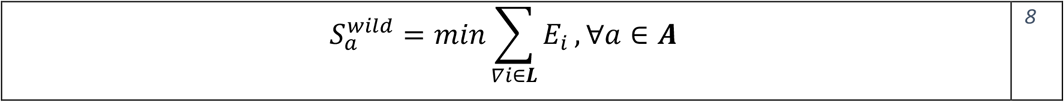

### Gene Essentiality Analysis

Let ***G*** represent the set of expressed genes. ∀*a* ∈ ***A***, we model each gene removal (*g* ∈ ***G***) sequentially to quantify the biological impact of its knockout for a given abstract and experimental dataset. The gene removal is basically imposed by forcing *E_g_* to be equal to zero (*E_g_* = 0) with *g* representing the gene that is being knocked out. Note that genes that appear in the pathway models and are not experimentally measured are considered as expressed and therefore we include them in the knockout process.

Afterwards the problem is solved (eq 8) and the minimum number of non-expressed genes active is calculated 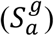. If 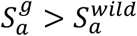 the gene is considered as essential for the cell to carry out that biological process (*a*) in the given pathway. Else, the gene is considered as not essential. In other words, if the new solution 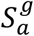 modelling the knockout of gene *g* requires the presence of more lowly expressed genes that the wild type 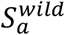, we assume that the removal of *g* is causing a significant impact to the phenotype represented by the gene expression dataset. The flow diagram corresponding to the methodology is summarised in **Figure 2**.

**Figure 2 -.**
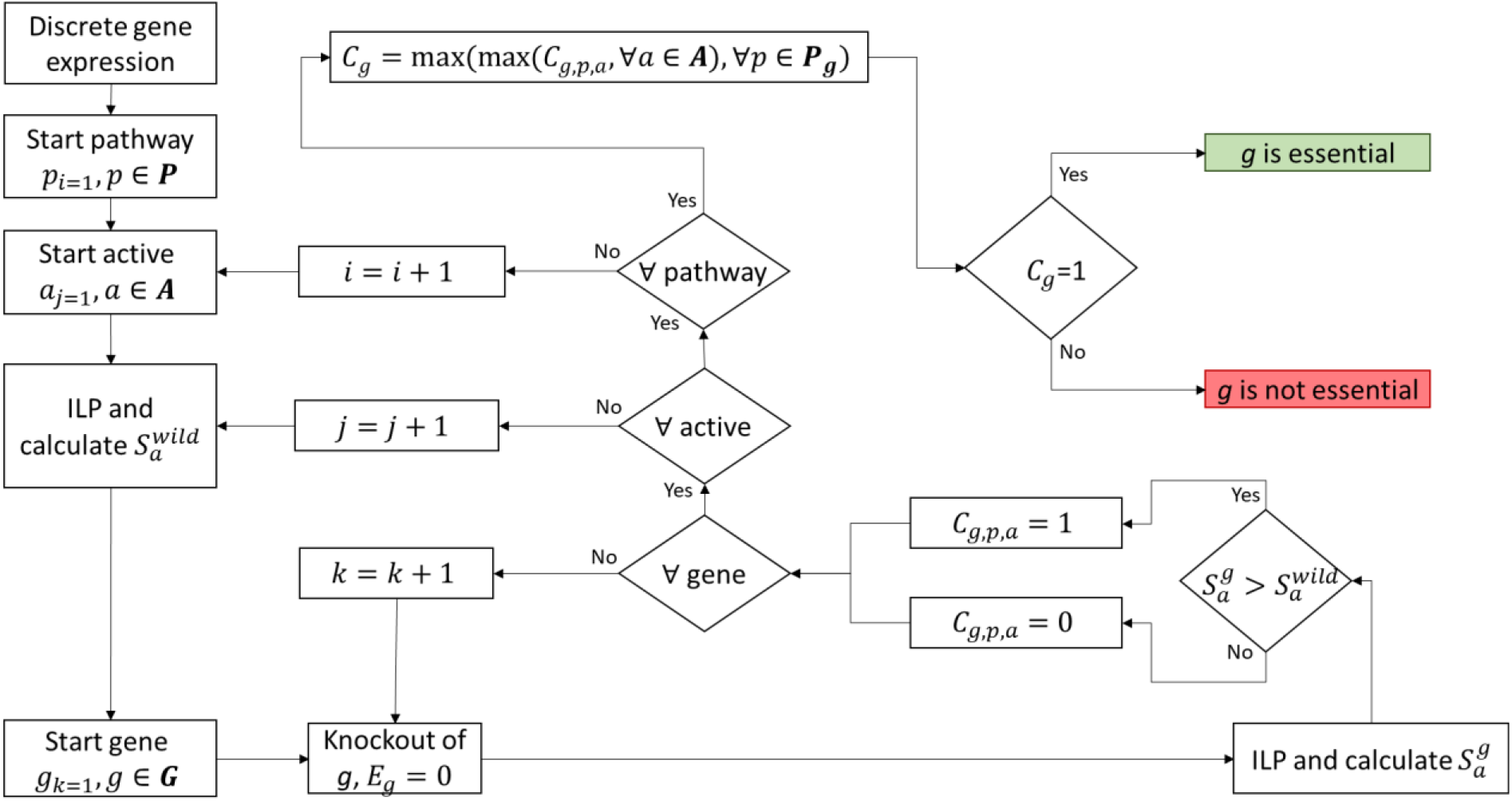
Flow diagram of the methodology. Starting from a specific experimental picture (discrete gene expression), we calculate the minimum number of non-expressed genes required to be active for the cell to sustain cellular life 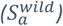. Then, we systematically knock-out one by one all the expressed genes g present in the pathway **P** (E_g_ = 0) and recalculate the minimum number of non-expressed genes required to be active for the cell to sustain cellular life 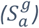. We define a gene as essential for a given active if 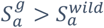. We repeat this process for all the genes, actives, and pathways included in the database. The essentiality of a gene g is finally defined as the maximum of all its essentiality predictions.

Note that the proposed methodology analyses every pathway independently and thus produces a prediction of essentiality for every gene and every active (gene complex or biological abstract) in the pathway. This means, that for a given gene, multiple predictions of essentiality can be produced in the same pathway (as many as there are elements in ***A*** for that pathway). Conceptually, our method assumes that if the gene is essential for at least one entity required to sustain cellular life, then its knockout would be fatal for the cell overall. Therefore, a gene is essential in a pathway, if it is essential for any of its actives.

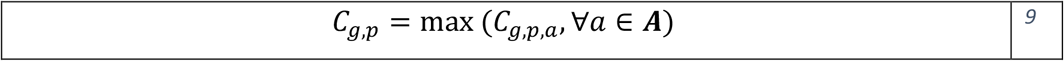

Where *C_g,p,a_* is a binary variable (*C_g,p,a_* ∈ {0,1}) that represents the essentiality of the gene *g* in the pathway *p* for the active *a*.

Moreover, different pathways are not completely disjoint sets and often have common genes. This means, that we can have more than one prediction of essentiality for a gene in different pathways. Similarly, we assume that if the gene is essential for at least one pathway, then its knockout would be fatal for the cell overall. Therefore, a gene is essential, if it is essential for any of its pathways.

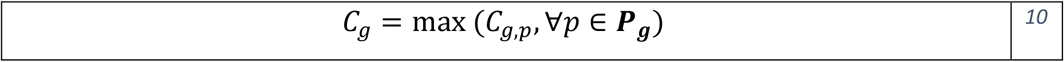

Where ***P_g_*** represents the set of pathways where the gene *g* is present.

#### Globally essential and globally not essential genes

If the knockout of a gene *g* leads to *C_g_* = 1 for each of the experimental datasets, the gene is considered globally essential. Similarly, if the knockout of a gene leads to *C_g_* = 0 for every single experimental dataset, the gene is considered globally not essential. Both globally essential and globally not essential genes are excluded from downstream analysis. Given the ulterior motives of the method, we are particularly interested in genes whose essentiality depends on the experimental dataset. Therefore, if a particular gene turns out to be essential in a cancer phenotype but not in the corresponding healthy tissue, we can identify it as a potential drug target.

#### Essentiality Congruity Score (ECS)

The proposed methodology assumes that predictions of essentiality (*C_g,p,a_* = 1) are more impactful than predictions of no essentiality (*C_g,p,a_* = 0) and the essentiality of a gene *g* is defined as the maximum of all its predictions (eqs 9 and 10). This assumption, however, is very susceptible to false positive predictions (not essential genes predicted as essential) that can have a huge impact on the obtained results. To address this issue, we defined the *Essentiality Congruity Score* (*ECS*) as:

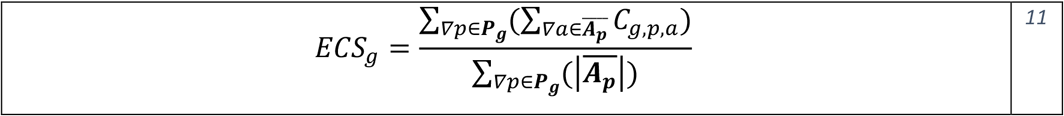

where *ECS_g_* is the Essentiality Congruity Score for the gene *g*, ***P_g_*** represents the set of pathways in which the gene *g* is present, 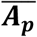 is the set of actives for the pathway *p* with at least one prediction of essentiality, and *C_g.p,a_* is the prediction of essentiality for the gene *g*, in the pathway *p*, and for the active *a*. *ECS_g_* = 0 means that none of the predictions for the gene *g* were of essentiality while *ECS_g_* = 1 means that 100% of the predictions were essential.

## Results

In this section, we show the results obtained with the proposed methodology when applied to a set of case scenarios: 1) a simple toy example showing the key conceptual aspects of the methodology and the functioning of the equations; 2) a case study using the gene essentiality data from the Achilles project illustrating the biological validity of the obtained results; 3) a breast cancer dataset and validate the obtained results in the literature.

### Toy Example

First, we considered a simplification of the *Wnt receptor signaling pathway, planar cell polarity pathway*, which is shown in **Figure 3** (Schaefer *et al*., 2009).The simplified subnetwork comprises four genes (WNT5A, FZD7, WNT3A and FCD1), two complexes (WNT5A/FZD7 and WNT3A/FZD1) and an abstract (Wnt receptor signaling pathway, planar cell polarity pathway). As mentioned earlier, the methodology comprises two main steps: (i) calculating the minimum number of non-expressed genes that we need to activate in order to trigger a given active *a* 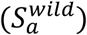 and (ii) performing an exhaustive *in-silico* gene knockout to find deletions that unavoidably lead to the need of activating extra non-expressed genes in order to trigger the given entity 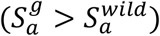.

**Figure 3 -.**
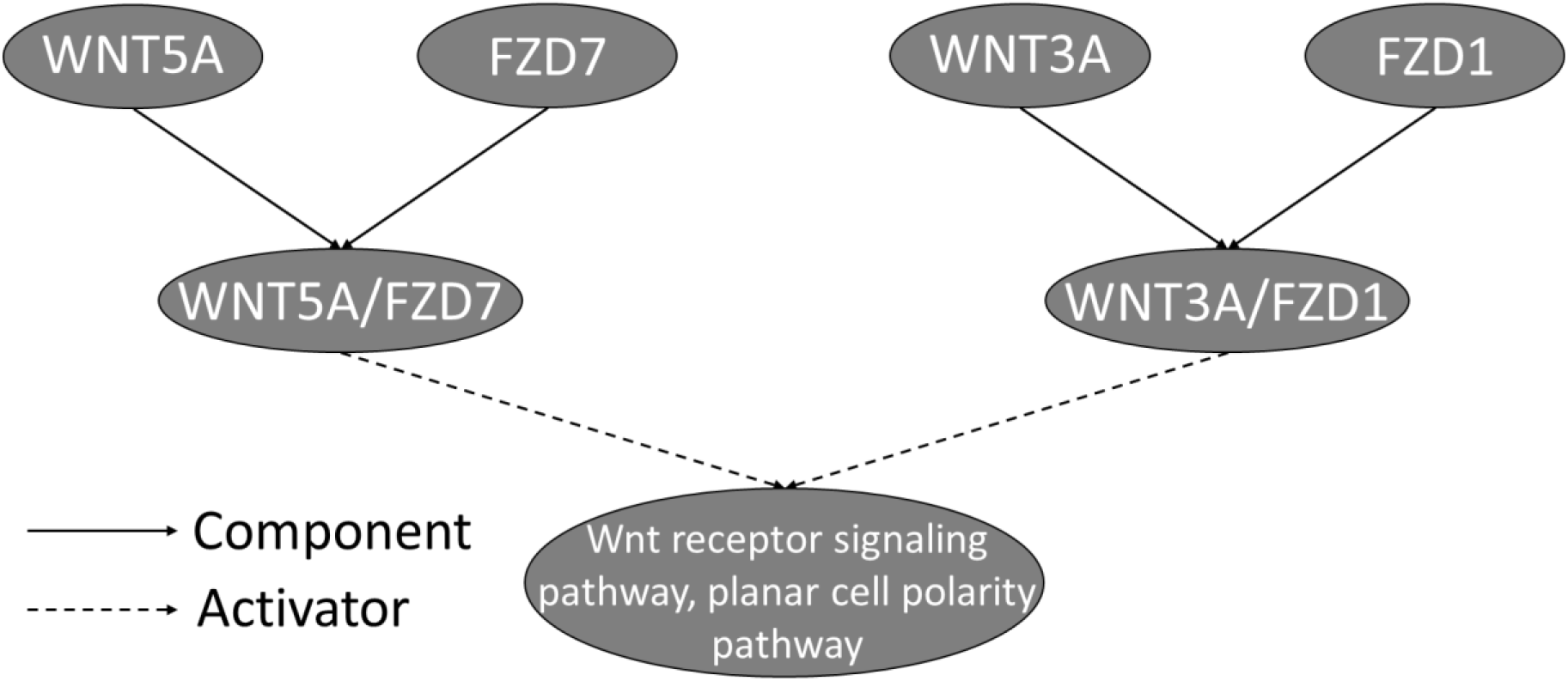
Toy Example. Graphical representation of the pathway activating the Wnt receptor signalling pathway, planar cell polarity pathway. Component-type interactions are represented with solid arrows whilst activation-type interactions are illustrated with dashed lines.

In the forthcoming lines we will define three scenarios based on simulated data. These scenarios show different solutions based on whether WNT5A and WNT3A are expressed or not while FZD7 and FZD1 are always expressed *i.e*., FZD7 & FZD1 ∈ **G**. **Table 1** summarises the solution of the different scenarios proposed. The complete solution of the mathematical model for each scenario is included in Supplementary Results 1.

**Table 1-.** Toy example solution. Possible scenarios when FZD7 and FZD1 are expressed. For each scenario, the expression values of each gene and the activity values of each entity are included. W5F7C represents the WNT5A/FZD7 complex, W3F1C represents the WNT3A/FZD1 complex, and ABSTR represents the Wnt receptor signalling pathway, planar cell polarity pathway. A) WNT5A and WNT3A are non-expressed. For the abstract to be active we need to activate one non-expressed gene (WNT3A in the example). B) WNT5A is not expressed and WNT3A is expressed. For the abstract to be active we do not need to activate any non-expressed gene. C) WNT5A is expressed and WNT3A is not expressed. For the abstract to be active we do not need to activate any non-expressed gene. B’) Scenario B after FZD7 is knocked-out. For the abstract to be active we do not need to activate any non-expressed gene. C’) Scenario C after FZD7 is knocked-out. For the abstract to be active we need to activate one non-expressed gene (WNT3A).

| Scenario | Gene Expression |  |  |  | Entity Activity |  |  |  |  |  |  |
| --- | --- | --- | --- | --- | --- | --- | --- | --- | --- | --- | --- |
|  | WNT5A | FZD7 | WNT3A | FZD1 | WNT5A | FZD7 | WNT3A | FZD1 | W5F7C | W3F1C | ABSTR |
| A | 0 | 1 | 0 | 1 | 0 | 0 | 1 | 1 | 0 | 1 | 1 |
| B | 0 | 1 | 1 | 1 | 0 | 0 | 1 | 1 | 0 | 1 | 1 |
| C | 1 | 1 | 0 | 1 | 1 | 1 | 0 | 0 | 1 | 0 | 1 |
| B' | 0 | 1 | 1 | 1 | 0 | 0 | 1 | 1 | 0 | 1 | 1 |
| C' | 1 | 1 | 0 | 1 | 0 | 0 | 1 | 1 | 0 | 1 | 1 |

#### (i) Minimum number of non-expressed genes required to activate an entity

The scenario shown in **Figure 4.A** has two expressed genes (FZD7 and FZD1) and two non-expressed genes (WNT5A and WNT3A). For the abstract to be active, one of the two complexes needs to be active. The condition for either complex is that both of its gene components need to be active. Thus, in scenario A we need to activate one non-expressed gene (WNT5A or WNT3A) for the abstract to be active 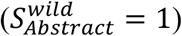. Scenarios B and C do not require the activation of any non-expressed gene to activate the abstract and therefore 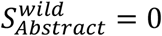 (for the detailed resolution, please refer to Supplementary Results 1).

**Figure 4 -.**
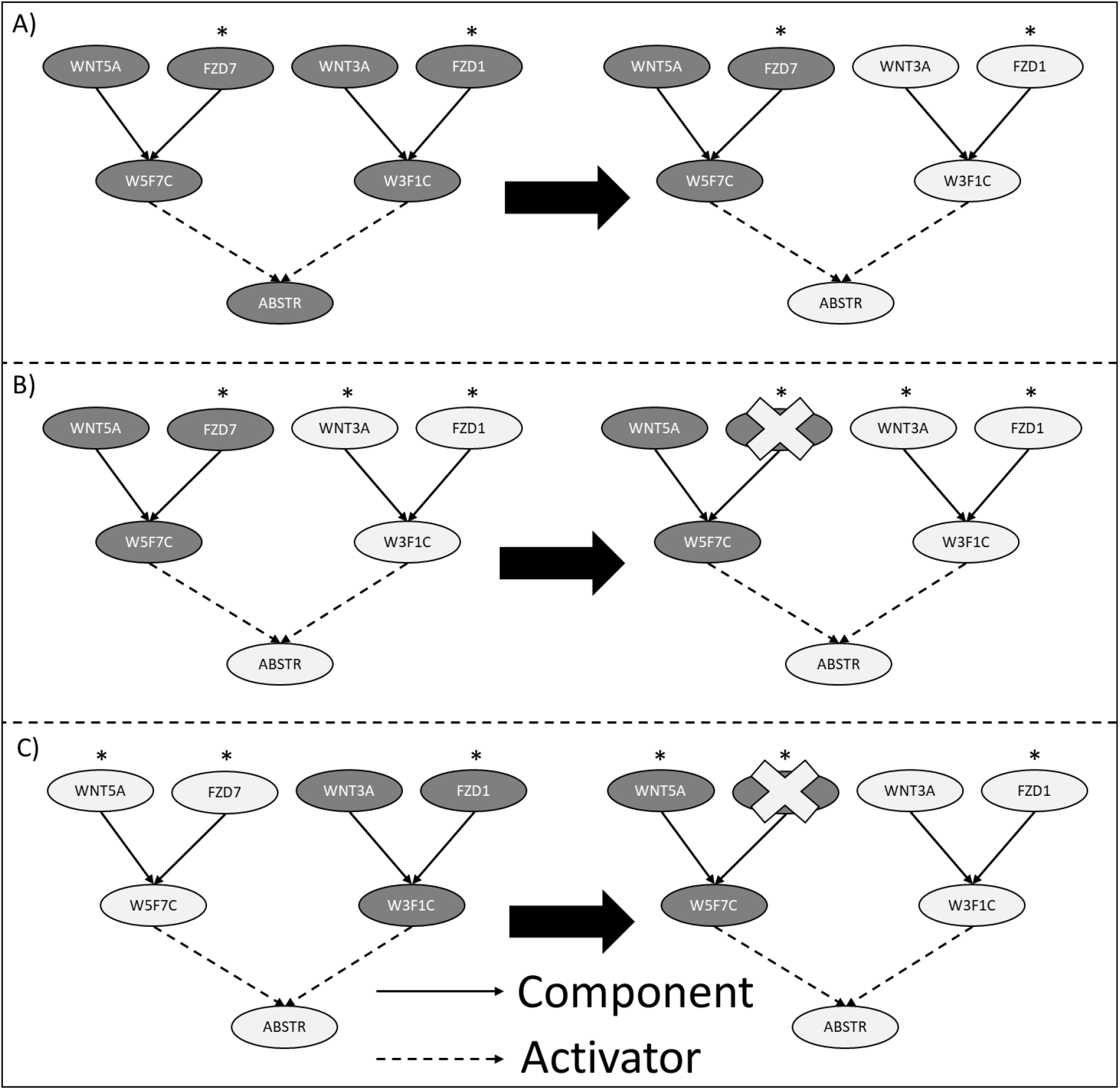
Toy example solution. Possible scenarios when FZD7 and FZD1 are expressed. W5F7C represents the WNT5A/FZD7 complex, W3F1C represents the WNT3A/FZD1 complex, and ABSTR represents the Wnt receptor signaling pathway, planar cell polarity pathway. Dark and ligth nodes represent inactive and active nodes in the final solution respectively, namely E_i_ = 0 and E_i_ = 1. The star on top of a gene g represents the expression state, in particular if g has a star it is expressed, namely g ∈ G, and not expressed otherwise, i.e. g ∈ L. A) WNT5A and WNT3A are not expressed. For the abstract to be active we need to activate one non-expressed gene (WNT3A in the example). B) WNT5A is not expressed and WNT3A is expressed. For the abstract to be active we do not need to activate any non-expressed gene. A knock-out of FZD7 does not require the activation of any non-expressed gene for the abstract to be active. C) WNT5A is expressed and WNT3A is not expressed. For the abstract to be active we do not need to activate any non-expressed gene. A knock-out of FZD7 requires the activation of one non-expressed gene (WNT3A) for the abstract to be active.

#### (ii) in-silico exhaustive gene knockout

In the scenario shown in **Figure 4.B**, a knock-out in FZD7 (*E_FZD7_* = 0) does not require the activation of any non-expressed gene because the abstract can be activated through the WNT3A/FZD1 complex and both its components are expressed, that is 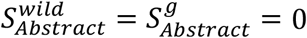. Therefore, in this scenario, FZD7 is not an essential gene. On the other hand, if we consider the scenario described in **Figure 4.C**, a knockout in FZD7 means that the WNT5A/FZD7 complex cannot be active, and thus the abstract needs to be activated via the WNT3A/FZD1 complex, which requires the activation of one non-expressed gene (WNT3A). In this scenario, 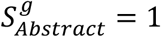, while 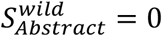 and therefore FZD7 is considered as an essential gene.

### Method Validation

To validate the biological relevance of the gene essentiality predictions of our method, for a given cell line, we compared the Achilles scores of the genes *g* predicted as essential (*C_g_* = 1) versus the scores of the genes predicted as not essential (*C_g_* = 0) (**Figure 5.A**). For the purpose of this analysis, globally essential (genes predicted as essential in all cell lines) and globally not essential genes (genes not essential in all the cell lines) were not included in the analysis (Methods section). This reduced the number of genes included in the comparison to 159.

**Figure 5 -.**
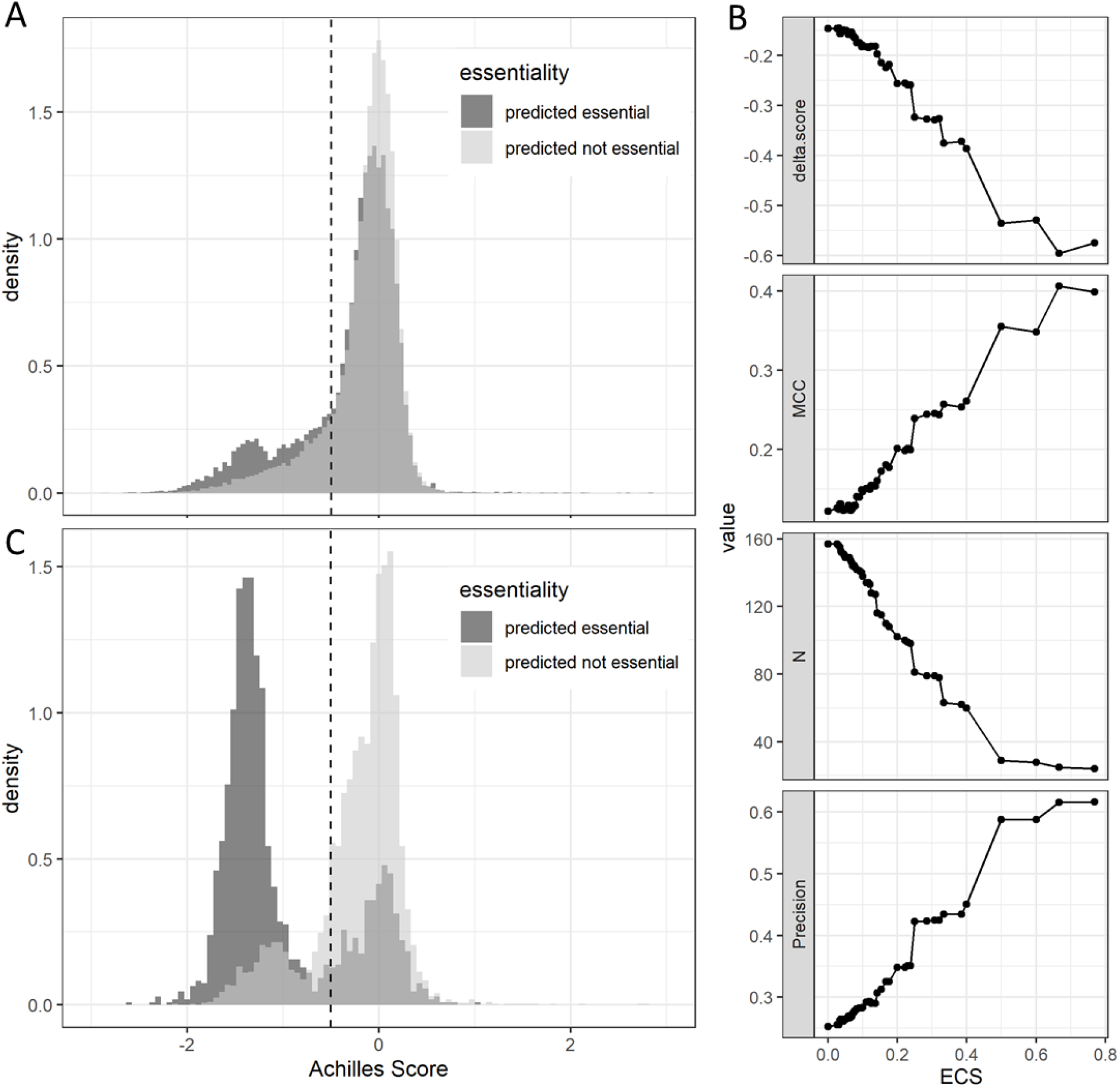
Method validation. A) Histogram showing the results from the validation of the method. The dark distribution shows the Achilles scores of those pair gene & cell-lines predicted as essential; the light distribution shows the Achilles scores of those predicted as not essential. Genes predicted as essential have significantly lower Achilles score than genes predicted as not essential (p-value = 6.4032^-246^). The average difference between both distributions is defined by the parameter delta.score =-0.1463. B) Impact of ECS in the performance of the method. Evolution of the results when different thresholds of ECS are used to define a gene as essential. delta.score: average difference in Achilles score between the genes predicted as essential and the genes predicted as not essential; MCC: Matthew’s Correlation Coefficient; N: number of genes included in the comparison; Precision: obtained precision assuming as real essential genes those with an Achilles score < −0.5. C) Histogram when MCC finds its maximum (ECS = 0.6667). The average difference in Achilles Score between genes predicted as essential and genes predicted as not essential becomes bigger (delta.score = −0.5954) and so does their significance (p-value = 0).

**Figure 5.A** shows how the genes predicted as essential have a significantly lower Achilles score than the genes predicted as not essential (p-value = 6.4032^-246^). The results illustrated in **Figure 5.A** follow the definition of essentiality represented in eqs 9 and 10 of the methods where a gene is considered essential if is predicted as essential for any active in any of the pathways where it appears. However, the ECS defined in eq 11 is a continuous score (*ECS* ∈ [0,1]) and allows to describe flexible threshold when defining the essentiality of a gene. For example, we can define genes as essential if their ECS is larger than a given threshold *th* (*C_g_* = 1, *if ECS_g_* > *th*). We studied the impact of applying different thresholds to the ECS by evaluating the evolution of the obtained results (**Figure 5.B**). For the purpose of this analysis, we defined a gene as really essential for a given cell-line if its Achilles score was below −0.5.

**Figure 5.B** shows how as the minimum ECS required to consider a gene as essential increases, so does the quality of the predictions. Most of the statistics shown in the different subfigures improve their performance when more demanding values of ECS are needed to define a gene as essential. We defined as the optimal cut-off the ECS where the MCC parameter finds its maximum (ECS = 0.67, MCC = 0.41, **Figure 5.C**). We selected the MCC because has been proven to be the optimal classifier for imbalanced data (Boughorbel et al, 2017). However, as the minimum required threshold increases, so does the number of genes considered globally not essential which decreases the number of genes included in the analysis (represented by N). **Figure 5.B** also shows the monotonically increasing behaviour of the Precision curve when the minimum required ECS to define essentiality increases. This is particularly interesting for reducing experimental validation costs, as we want to make sure that genes predicted as essential are indeed essential while genes predicted as not essential are not as relevant.

#### Synergistic behaviour of the method

This gene essentiality method finds its success on the synergy between three different factors: 1) biologically relevant gene expression data, 2) a robust prior-knowledge-network (PKN), and 3) the mathematical formulation described in the methods section. Alterations in each of these fundamental pillars affect downstream results increasing the number of false positive predictions. To test the first pillar, biologically relevant gene expression data, we fed the method with “nonsense” expression data by inverting the binary scores obtained from The Gene Expression Barcode 3.0 (McCall et al., 2012), (McCall et al., 2014). This, reduced the maximum MCC (starting from a baseline of 0.41 using a ECS of 0.67) to 0.1 (using a ECS of 0.5). To validate the need of a representative prior-knowledge network we repeated the analysis using only the subset of 50 NCI-PID pathways that were labelled as tumorigenic which increased the maximum MCC to 0.53 (using a ECS of 0.67). Finally, we evaluated that this improvement in MCC was only present when the gene expression data was biologically meaningful. To that end, we repeated the analysis using the subset of NCI-PID pathways and the “nonsense” expression data obtaining a MCC of 0.12 (using a ECS of 0.9). The reader should refer to Supplementary Results 2 for the complete evaluation. When compared with other state of the art methods (Cubuk et al., 2018) our method produces less false positives (Supplementary Results 2).

### Case Study – Breast Cancer

Finally, we applied the gene essentiality method to Breast Cancer patient samples (Maubant S, 2012), (Maire V, 2013) and looked for genes significantly predicted as essential in cancer patients using hypergeometric tests. For this purpose, technical duplicates were considered as independent samples. A gene was considered essential for a given patient if ECS > 0. The same procedure was repeated for the different cancer subtypes.

**Table 2** shows the top 10 results for the Healthy vs BRCA case while Supplementary Table 2 shows the top 10 results for the group-specific comparison. The complete lists can be found in Supplementary Tables 3 and 4. In the following lines, we will highlight the relevance of the top 4 (elbow criterion) genes reported in **Table 2** with a higher coverage of patients by relying on existing knowledge in the literature.

**Table 2 -.** BRCA essentiality results. q: number of cancer patients predicted as essential, k: number of total patients, including healthy and cancer, predicted as essential. Number of cancer samples = 153, Total number healthy samples = 11. Score: average Achilles score across all cell lines.

| geneID | p.hyper | p.adj | q | k | score |
| --- | --- | --- | --- | --- | --- |
| **RACGAP1** | 3.25E-07 | 4.91E-05 | 118 | 118 | -1.45764 |
| **MIB1** | 2.33E-06 | 1.76e-04 | 110 | 110 | -0.34808 |
| **EZR** | 1.02E-05 | 5.13e-04 | 103 | 103 | 0.094074 |
| **PCNA** | 1.94E-05 | 7.34e-04 | 126 | 128 | -1.88689 |
| TUBG1 | 1.40e-04 | 4.21e-03 | 88 | 88 | -1.31599 |
| CASP3 | 5.20e-04 | 9.81e-03 | 79 | 79 | 0.073582 |
| PKCDELTA | 5.20e-04 | 9.81e-03 | 79 | 79 | NA |
| SDC2 | 5.20e-04 | 9.81e-03 | 79 | 79 | -0.06373 |
| **BIRC3** | 1.68e-03 | 2.31e-02 | 89 | 90 | 0.149018 |
| GNAI1 | 1.68e-03 | 2.31e-02 | 70 | 70 | -0.10703 |

### RACGAP1: Rac GTPase-activating protein 1

RACGAP1 is a protein involved in several biological processes including cell cycle, cell division, and differentiation and with a key role in various cellular phenomena including cytokinesis, invasive migration and metastasis. Increased expression of RACGAP1 protein has been previously associated with poor survival as well as significantly associated with increased tumour malignancy in colorectal cancer (Imaoka et al, 2015). It has been shown that its knockdown – in combination with radiotherapy – is associated with a decrease of tumour viability and invasiveness in 4T1 mouse models (Wu et al, 2019).

### PCNA: Proliferating cell nuclear antigen

PNCA is a protein involved in DNA replication by increasing the processivity of DNA polymerase delta. Immunohistochemical staining of PCNA has been used extensively in breast cancer diagnosis and prognosis (Malkas et al, 2006). It has been shown that targeting the EGFR/PCNA signalling suppresses tumour growth of triple-negative breast cancer cells (Yu et al, 2013) and inhibit cancer growth in neuroblastoma and breast cancer mouse xenograft models (Choe et al, 2017).

### MIB1: Mindbomb E3 ubiquitin protein ligase 1

MIB1 is a protein that positively regulates Notch signaling by ubiquitinating the Notch receptors, thereby facilitating their endocytosis. It has been shown that MicroRNA-198 suppresses prostate tumorigenesis by targeting MIB1 (Ray et al, 2019). MicroRNA-198 also represses cell proliferation and migration and promotes cell adhesion in breast cancer cells (Hu et al 2017).

### EZR: Ezrin

EZR is protein that plays a key role in cell surface structure adhesion, migration and organization. Its inhibition synergizes with lapatinib in a PKC-dependent fashion to inhibit proliferation and promote apoptosis in HER2-positive breast cancer cells (Jeong et al, 2019). EZR inhibition in hepatocellular carcinoma (HCC) cells decreases their migratory and invasive potential (Zhang et al, 2006).

## Discussion

This article introduces a new methodology for the *in-silico* identification of essential genes which integrates high-throughput gene expression data with predefined biological pathways to provide patient-specific gene essentiality predictions. This method uses a mathematical formulation that identifies the number of non-expressed genes required to be active for the cell to sustain life, here modelled by the activation of a relevant biological task. This work expands the ideas behind existing CBM-based methodologies going beyond metabolism by considering multisystem networks (Schaefer et al., 2009).

We have validated the proposed methodology using a set of 452 cancer cell lines derived from the Cancer Cell Line Encyclopedia where the essential genes had been previously identified using CRISPR knockouts (Achilles Project). When compared to competing methods, our approach identifies essential genes with fewer false positives. Because cell-lines do not represent the entire complexity of cancer, we have further supported the obtained essential genes in an independent breast cancer dataset using existing literature.

The mathematical formulation presented in the methods section makes it possible to have several predictions of essentiality for the same gene. Due to the nature of the problem, initially a single prediction of essentiality was a sufficient condition to consider the gene as essential thus these multiple predictions were summarized into their maximum for each gene. This summarization is very susceptible to false positive results which can have a huge impact in downstream results. We have shown how the integration of multiple predictions into the *Essentiality Congruity Score* (ECS) improves our ability to identify essential genes.

The presented methodology finds its success on the synergy between its three core constituents: biologically relevant gene expression data, a robust prior-knowledge-network that effectively captures cancer biological events and the constraint-based mathematical model described in the methods section.

We have demonstrated that all three elements are necessary by modifying individual constituents. We have proven that missense input data (produced by inverting the discrete expression values) does not yield to valid results. We have also shown that including pathways that do not represent tumorigenic events worsen the essentiality predictions. Finally, we have proven how diluting the impact of false positive predictions derived from the methodology using the ECS further improves the precision when identifying essential genes.

The mathematical formulation described in the methods section distinguishes between expressed genes and non-expressed genes. This discrimination, however, is derived from continuous gene expression data, which was previously discretized using The Gene Expression Barcode 3.0 (McCall et al, 2012), (McCall et al, 2014). This work does not directly tackle this issue but the selection of discretization strategy can have a tremendous impact on downstream results.

The present methodology assumes that all the actives (abstracts + complexes) included in the PKN are equally relevant for the cell to sustain life. This represents an oversimplification of the reality as not all the actives will affect the cell in the same way. We have shown that removing pathways that do not capture tumorigenic events improve the obtained results demonstrating that there needs to exist harmony between the biological network and the mathematical model.

The advent of *in-silico* approaches predicting essential genes will pave the way for precision medicine by identifying potential drug targets whose deletion can induce death in tumour cells (Tsherniak *et al*., 2017). The work presented here contributes in this direction. However, further efforts are required in order to develop disruptive *in-silico* methodologies that accounts for further biophysical knowledge, such as dynamic models or multi-omics data. Overcoming this ambitious challenge will set the foundations for addressing biological questions that were unreachable before.

## Acknowledgements

The authors would like to thank Matthew Trotter and Manuel Sanchez Castillo for the fruitful discussions and the useful suggestions and comments.

## Supplementary Results

### Supplementary Results 1: Toy Example – Solution of the mathematical model

In the forthcoming lines, we will solve the mathematical model for different scenarios of the toy example included in **Figure 4**. The different scenarios involve different genes expressed and not expressed. **Table 1** summarises the expression of all the genes and the activity of all the entities included in the pathway for each scenario.

As mentioned in the methods, the methodology comprises two main steps: (i) calculating the minimum number of non-expressed genes that we need to activate in order to activate a given active 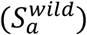 and (ii) performing an *in-silico* exhaustive gene knockout to find deletions that unavoidably lead to the need of activating extra non-expressed genes in order to activate the active 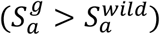. Scenario A will be used to illustrate how the minimum number of non-expressed expressed genes needed active is calculated. Scenarios B and C will be used to illustrate how essential genes are obtained.

#### Scenario A

In this scenario, FZD7 and FZD1 are expressed (*G* = {FZD7, FZD1}) while WNT5A and WNT3A are not expressed (*L* = {*WNT5A, WNT3A*}). The Wnt receptor signalling pathway, planar cell polarity pathway - from now on, the abstract – is the entity required for the cell to sustain life, that is, ***A*** = {*Abstract*}. Subject to the restriction modelled by eq 7, any valid solution needs for the abstract to be active:

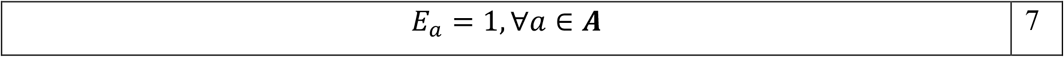

that is, *E_Abstract_* = 1.

The parents of the abstract are two complexes (WNT5A/FZD7, and WNT3A/FZD1) that are connected to the abstract via activation interactions. According to the rules describing the activators presented in the methods section, the activation of the abstract is determined by its activation/inhibition state, which according to eq 5, is defined as:

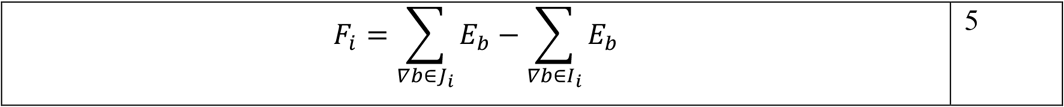

In this case, the abstract (*i*) does not have any inhibitors (*I_Abstract_* = ø) and has two activators, that is, *J_Abstract_* = {WNT5A/FZD7, WNT3A/FZD1}. Therefore:

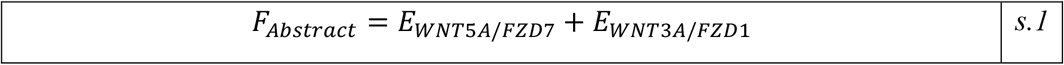

According to the equations for activators and inhibitors, the activation of the abstract is defined as:

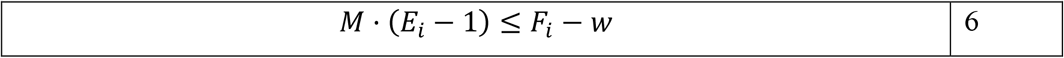

Hence, if we substitute eq s.1 in eq 6 with the standard parameters (M = 1000, w = ½), and bearing in mind that *E_Abstract_* = 1, we obtain:

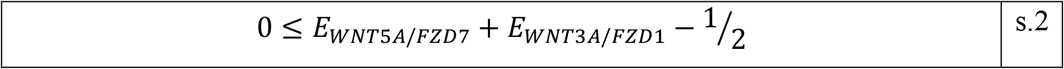

If we clean up the expression above,

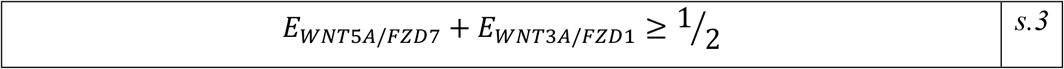

*E_WNT5A/FZD7_* and *E_WNT3A/FZD1_* are binary variables (*E_WNT5A/FZD7_, E_WNT3A/FZD1_* ∈ {0,1}) thus, for the first expression to be true, at least one of the two complexes needs to be active. The reader should note that the solution where both complexes are active (*E_WNT5A/FZD7_* = *E_WNT3A/FZD1_* = 1) is also feasible. In this example, we will solve the equations for the WNT5A/FZD7 complex (*E_WNT5A/FZD7_* = 1) but the reader should know that the same logic can be applied for the WNT3A/FZD1 complex.

The WNT5A/FZD7 complex has two parents that are connected to the child via component interactions. Following the rules for components introduced in the methods section (eqs 1 and 2) the activity of a complex is defined as:

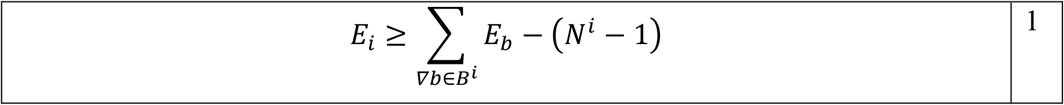

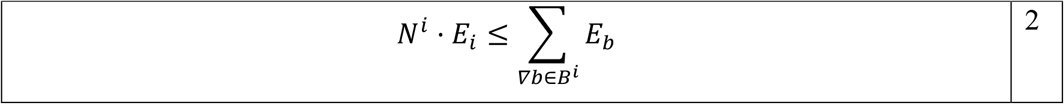

In this case, *B^WNT5A/FZD7^* = {WNT5A, FZD7} and *N^WNT5A/FZD7^* = 2. Therefore:

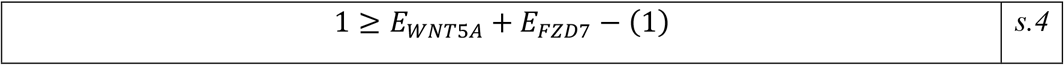

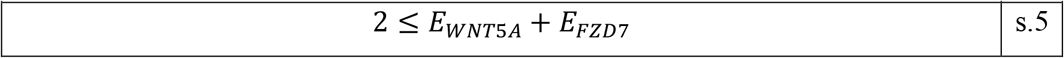

If we clean up the expressions above,

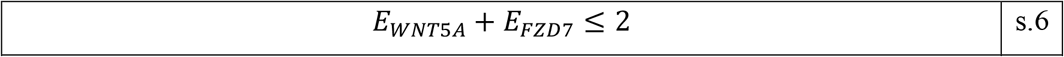

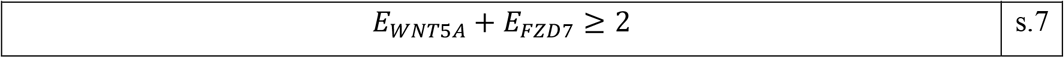

Thus,

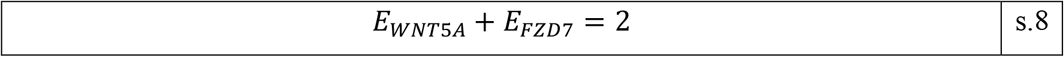

*E_WNT5A_* and *E_FZD7_* are binary variables (*E_WNT5A_, E_FZD7_* ∈ {0,1}), thus this equation is only true when *E_WNT5A_* = *E_FZD7_* = 1, that is, both component genes have to be active. As we can see in **Figure 4.A**, FZD7 is expressed but WNT5A is not expressed. Therefore, for the abstract to be active, we need at least one non-expressed gene active (WNT5A). Thus, in this scenario we have three feasible solutions:

- Solution 1: Activate WNT5A (*E_WNT5A_* = 1).
- Solution 2: Activate WNT3A (*E_WNT3A_* = 1).
- Solution 3: Activate WNT5A and WNT3A (*E_WNT5A_* = *E_WNT3A_* = 1).

However, not all three solutions are equally good as the objective function of the model defines as the optimal solution the one that minimizes the number of lowly expressed entities active in the final solution (eq 8):

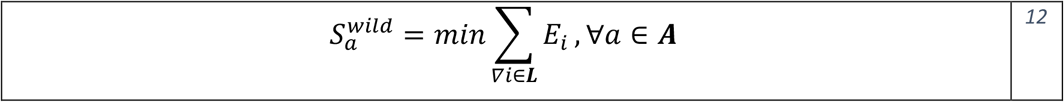

Solution 3 requires two non-expressed genes to be active 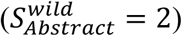, thus is a suboptimal solution. Solutions 1 and 2 require one non-expressed gene to be active, thus they are optimal solutions 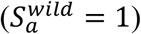.

#### Scenario B

In this scenario, FZD7, WNT3A, and FZD1 are expressed (*G* = {*FZD7, WNT3A, FZD1*}) while WNT5A is not expressed (*L* = {*WNT5A*}). Given that the topology of the pathway is the same, the solution of the mathematical model is analogous to scenario 1 until the calculation of the objective function. In this case, the number of feasible solutions is two:

- Solution 1: Do not activate any non-expressed genes.
- Solution 2: Activate WNT5A (*E_WNT5A_* = 1).

The first solution is the optimal solution as it does not require any non-expressed gene to be active, that is, 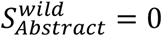).

If we knock-out FZD7 (*E_FZD7_* = 0), the solution that requires WNT5A to be active (Solution 2) is not feasible anymore because following the rules for components introduced in the methods section, the activity of the WNT5A/FZD7 complex is defined as reflected in eq s.8, that is, *E_WNT5A_* + *E_FZD7_* = 2

*E_WNT5A_* and *E_FZD7_* are binary variables (*E_WNT5A_*, *E_FZD7_* ∈ {0,1}), thus this equation is only true when *E_WNT5A_* = *E_FZD7_* = 1, that is, both component genes have to be active. This time, however, we have knocked-out FZD7 (*E_FZD7_* = 0) so this equation will never be true. That is, a knock-out of FZD7 leaves us with one feasible solution:

- Solution 1: Do not activate any non-expressed genes.

The only feasible solution is, of course, the optimal and it does not require any non-expressed gene to be active, that is, 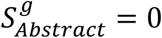. Note how 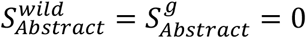. Therefore, in Scenario B, FZD7 is not an essential gene.

#### Scenario C

In this scenario, WNT5A, FZD7, and FZD1 are expressed (*G* = {*WNT5A, FZD7, FZD1*}) while WNT3A is not expressed (*L* = {*WNT3A*}). Given that the topology of the pathway is the same, the solution of the mathematical model is analogous to scenario A until the calculation of the objective function. In this case, the number of feasible solutions is two:

- Solution 1: Do not activate any non-expressed genes.
- Solution 2: Activate WNT3A (*E_WNT3A_* = 1).

The first solution is the optimal solution as it does not require any lowly expressed gene to be activated, that is, 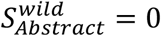.

If we knock-out FZD7 (*E_FZD7_* = 0), the solution that does not require any non-expressed genes active (Solution 1) is not feasible anymore. The demonstration has already been shown for Scenario B. This time, the only feasible solution after a knock-out of FZD7 is:

- Solution 2: Activate WNT3A (*E_WNT3A_* = 1).

The only feasible solution is, of course, the optimal and it requires one non-expressed gene active, that is, 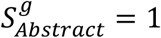. Note that his time 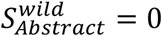 while 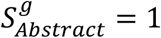. Therefore, in Scenario C, FZD7 is an essential gene.

### Boolean model

In supplementary section 1, we have seen that for the abstract to be active, at least one of the two gene complexes needs to be active. This intrinsically represents a Boolean equation that can be represented as:

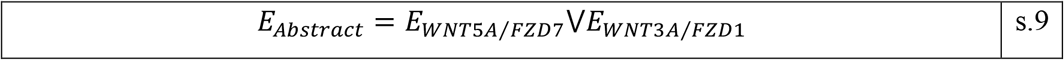

Where V represents the binary operator “or” and *abstract* refers to the Wnt receptor signaling pathway, planar cell polarity pathway.

On the other hand, for either complex to be active, both its component genes need to be active. This can also be represented following Boolean rules as:

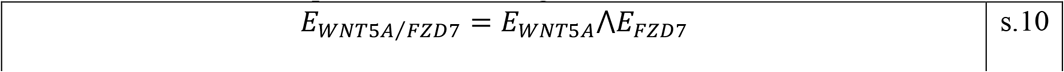

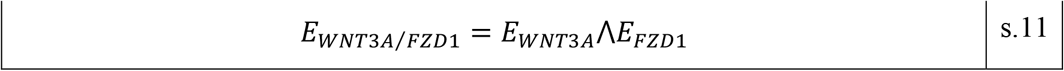

Where Λ represents the binary operator “and”.

When substituting eqs s.10 and s.11 in eq s.9, the activity of the abstract is directly expressed in terms of the activity of the genes:

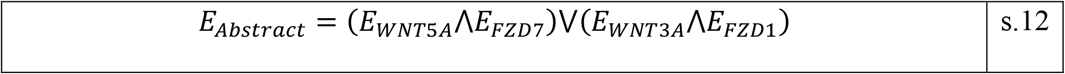

It should be noted that ILP eq 8 is fundamentally representing a set of Boolean expressions. However, when modelling a standard network, the emerging Boolean rules are too intricate and the use of ILP is required in order to address the problem. This growth in complexity emerges when incrementing the number of nodes and vertices in the network or when feedback loops are included.

### Supplementary Results 2: Fundamental pillars of the method

This gene essentiality method finds its success on the synergy between three different factors: biologically relevant gene expression data, a robust prior-knowledge-network (PKN), and the mathematical formulation described in the methods section. The following lines study how alternations in each of these fundamental pillars affect downstream results and how indeed we need all three of them to produce valid results.

#### Analysis of gene-expression data

First, we studied the need of meaningful gene expression data. To that end, we generated a “nonsense” expression dataset by inverting the binary scores obtained from The Gene Expression Barcode 3.0 (McCall et al., 2012), (McCall et al., 2014) in the CCLE dataset, that is, substituting the 1s (expressed genes) for 0s (non-expressed genes) and vice-versa (1 → 0,0 → 1), and repeated the whole analysis for this newly generated dataset. These results are summarized in **Error! Reference source not found.**. **Error! Reference source not found.** shows how for practically any ECS threshold, the dataset with the original expression profile data (original) produces better results than the nonsense dataset (inverse data). This clearly shows how gene expression data drives the obtained gene essentiality predictions and that these are not entirely the result of the network topology.

#### Analysis of prior-knowledge-network

Second, we studied the need of a representative prior-knowledge-network. In the validation dataset, we are comparing Achilles scores between genes predicted as essential and genes predicted as not essential in different of cell lines. These cell lines come from the Cancer Cell Line Encyclopedia, therefore essential means essential for cancer. However, not all the pathways introduced in the NCI-PID database are equally relevant for cancer biology, not all of them represent hazardous effects that can be avoided by knocking-out essential genes.

For example, the “Hypoxia-inducible factor (HIF)-2 alpha” is a hazardous pathway strongly correlated with cancer (Leek et al, 2002). On the other hand, the “LKB1 signaling events” is a tumour suppressor pathway (Momcilovic & Shackelford, 2015). The results for the predictions of essentiality for each of these pathways are summarized in **Table s.1**. **Table s.1** shows how the predictions for the tumorigenic pathway are superior to those of the tumor suppressor pathway. This agrees with the fact that the Achilles scores used in the validation represent essentiality for cancer cell lines. Therefore, it is expected that a tumor suppressor pathway is not going to be able to properly capture this essentiality.

**Table s.1 -.** Pathway validation. Performance of the gene essentiality predictions in two different pathways: Hypoxia-inducible factor (HIF)-2 alpha (HIF-2 A), and LKB1 signaling events (LKB1). The HIF-2 A pathway is a tumorigenic pathway while the LKB1 pathway is a tumor suppressor pathway.

| Pathway | N | TP | FP | FN | TN | MCC |
| --- | --- | --- | --- | --- | --- | --- |
| HIF-2 A | 956 | 476 | 0 | 4 | 476 | 0.9917 |
| LKB1 | 1912 | 2 | 459 | 44 | 1407 | -0.0725 |

In order to evaluate the impact of the PKN on the obtained ECS, we manually annotated (AA & JP, double blinded) the NCI-PID pathways in three different categories: tumorigenic (category 1), tumor suppressor (category 2), and unrelated with cancer (category 3). The complete annotated list is included in **Supplementary Table 1**. We performed these annotations for the 84 pathways that were included in the analysis. That is, the 92 pathways that did not have any prediction of essentiality for any cell-line included in the validation dataset (static pathways) were excluded from the analysis and thus, the annotation. 50 of these 84 pathways were annotated as tumorigenic (category 1). We hypothesized that including only tumorigenic pathways in the validation dataset would improve the results. Results from this analysis are summarised in **Figure s.1**. **Figure s.1** shows how indeed including only tumorigenic pathways further improves the predictive ability of our method, particularly when the minimum ECS required to define essentiality is high.

**Table s.2 -.**
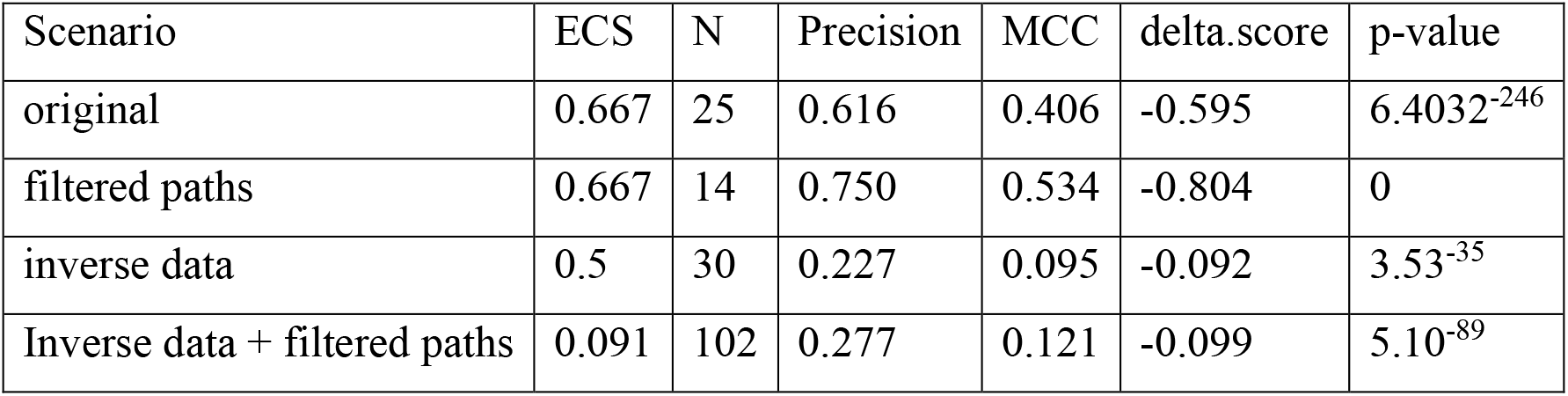
Optimal result for each model. Optimality was defined as the threshold were MCC was maximum. ECS: minimum ECS required to define essentiality. N: number of genes included after removing globally essential and globally not essential genes. Precision: Precision. MCC: Matthew’s Correlation Coefficient. delta.score: average difference in Achilles score between genes predicted as essential and genes predicted as not essential. p-value (t-test): obtained p-value from a t-test comparing the Achilles score of the genes predicted as essential versus the genes predicted as not essential (one-tailed).

| Scenario | ECS | N | Precision | MCC | delta.score | p-value |
| --- | --- | --- | --- | --- | --- | --- |
| original | 0.667 | 25 | 0.616 | 0.406 | -0.595 | 6.4032<sup>-246</sup> |
| filtered paths | 0.667 | 14 | 0.750 | 0.534 | -0.804 | 0 |
| inverse data | 0.5 | 30 | 0.227 | 0.095 | -0.092 | 3.53<sup>-35</sup> |
| Inverse data + filtered paths | 0.091 | 102 | 0.277 | 0.121 | -0.099 | 5.10<sup>-89</sup> |

For completeness, we also ran this analysis using the nonsense dataset (inverted data + filtered pathways) and saw how selecting only tumorigenic pathways does not improve the results when these come from meaningless biological data (**Figure s.1**). In each case, we selected the optimal required ECS where the MCC parameter finds its maximum. **Table s.2** summarises the results derived from the optimal threshold in each case. We selected the MCC because has been proven to be the optimal classifier for imbalanced data (Boughorbel et al, 2017).

**Figure s.1 -.**
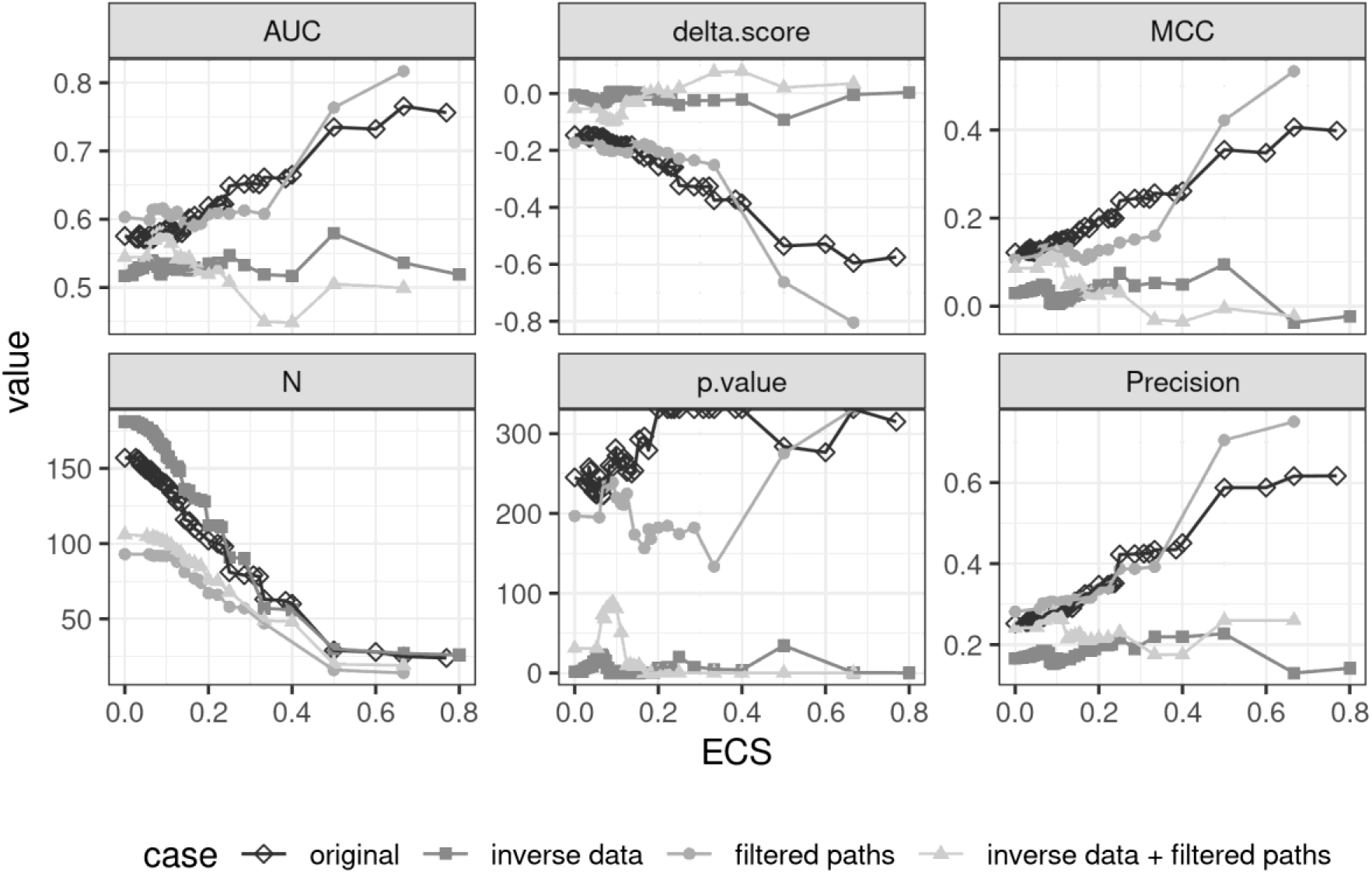
Evaluation of fundamental pillars of the methodology. Four different models were tested. Original: valid gene expression data and complete set of pathways from NCI-PID; inverse data: nonsense gene expression data (inverted) and complete set of pathways; filtered paths: valid gene expression data and tumorigenic pathways; and inverse data + filtered paths: nonsense gene expression data (inversed) and tumorigenic pathways.

These results further demonstrate the need of all three fundamental pillars in order to produce valid results: biologically meaningful gene expression data, a robust prior-knowledge-network (PKN), and a mathematical formulation valid for the identification of essential genes.

#### Comparative with other state of the art methods

Then, we compared the essentiality predictions obtained using our method with those obtained by another method available in the literature for the identification of essential genes (metabolizer, Cubuk et al, 2018). To this end, we compared the Achilles score of those genes predicted as essential by our methodology (original implementation) with the Achilles score of those genes predicted as essential by metabolizer (t-test). This analysis revealed that the genes predicted as essential by our method have significantly lower Achilles score than those predicted as essential by metabolizer (p.value = 7.7376e-37, delta = −0.2073).

---

**Supplementary Table 1 -.** Pathway annotation.

| source | id | Path | score |
| --- | --- | --- | --- |
| NCI-PID | 36875 | Cellular roles of Anthrax toxin | 2 |
| NCI-PID | 36879 | TCR signaling in naive CD8+ T cells | 3 |
| NCI-PID | 36882 | Nectin adhesion pathway | 1 |
| NCI-PID | 36892 | Aurora B signaling | 1 |
| NCI-PID | 36894 | RAC1 signaling pathway | 1 |
| NCI-PID | 36896 | IL2 signaling events mediated by PI3K | 1 |
| NCI-PID | 36899 | ErbB1 downstream signaling | 1 |
| NCI-PID | 36902 | TGF-beta receptor signaling | 1 |
| NCI-PID | 36905 | PLK2 and PLK4 events | 1 |
| NCI-PID | 36906 | Stabilization and expansion of the E-cadherin adherens junction | 1 |
| NCI-PID | 36909 | Signaling events mediated by HDAC Class I | 1 |
| NCI-PID | 36910 | Regulation of Notch signaling | 1 |
| NCI-PID | 36911 | Regulation of Androgen receptor activity | 2 |
| NCI-PID | 36912 | E-cadherin signaling in the nascent adherens junction | 1 |
| NCI-PID | 36915 | ATF-2 transcription factor network | 1 |
| NCI-PID | 36920 | Osteopontin-mediated events | 1 |
| NCI-PID | 36922 | Signaling events regulated by Ret tyrosine kinase | 1 |
| NCI-PID | 36927 | FoxO family signaling | 2 |
| NCI-PID | 36929 | Presenilin action in Notch and Wnt signaling | 1 |
| NCI-PID | 36930 | Reelin signaling pathway | 2 |
| NCI-PID | 36932 | BARD1 signaling events | 2 |
| NCI-PID | 36935 | Signaling events mediated by PRL | 1 |
| NCI-PID | 36937 | Class I PI3K signaling events mediated by Akt | 1 |
| NCI-PID | 36938 | Signaling events mediated by focal adhesion kinase | 1 |
| NCI-PID | 36939 | Atypical NF-kappaB pathway | 1 |
| NCI-PID | 36940 | Retinoic acid receptors-mediated signaling | 2 |
| NCI-PID | 36941 | EGF receptor (ErbB1) signaling pathway | 1 |
| NCI-PID | 36947 | Signaling events mediated by VEGFR1 and VEGFR2 | 2 |
| NCI-PID | 36950 | Nephrin/Neph1 signaling in the kidney podocyte | 2 |
| NCI-PID | 36951 | Glypican 3 network | 2 |
| NCI-PID | 36952 | amb2 Integrin signaling | 3 |
| NCI-PID | 36956 | Regulation of cytoplasmic and nuclear SMAD2/3 signaling | 1 |
| NCI-PID | 36959 | mTOR signaling pathway | 1 |
| NCI-PID | 36962 | PDGFR-beta signaling pathway | 1 |
| NCI-PID | 36964 | Angiopoietin receptor Tie2-mediated signaling | 2 |
| NCI-PID | 36967 | p75(NTR)-mediated signaling | 3 |
| NCI-PID | 36968 | E-cadherin signaling in keratinocytes | 2 |
| NCI-PID | 36969 | ErbB2/ErbB3 signaling events | 1 |
| NCI-PID | 36970 | Role of Calcineurin-dependent NFAT signaling in lymphocytes | 2 |
| NCI-PID | 36974 | IFN-gamma pathway | 2 |
| NCI-PID | 36976 | CXCR4-mediated signaling events | 1 |
| NCI-PID | 36977 | Signaling events mediated by Stem cell factor receptor (c-Kit) | 1 |
| NCI-PID | 36978 | IL4-mediated signaling events | 3 |
| NCI-PID | 36981 | Caspase cascade in apoptosis | 2 |
| NCI-PID | 36982 | PDGFR-alpha signaling pathway | 1 |
| NCI-PID | 36984 | Internalization of ErbB1 | 1 |
| NCI-PID | 36985 | C-MYB transcription factor network | 1 |
| NCI-PID | 36986 | CD40/CD40L signaling | 1 |
| NCI-PID | 36989 | a6b1 and a6b4 Integrin signaling | 2 |
| NCI-PID | 36994 | EPHA2 forward signaling | 1 |
| NCI-PID | 36995 | HIF-2-alpha transcription factor network | 1 |
| NCI-PID | 36996 | LPA receptor mediated events | 1 |
| NCI-PID | 37001 | IL2 signaling events mediated by STAT5 | 1 |
| NCI-PID | 37002 | LKB1 signaling events | 3 |
| NCI-PID | 37004 | CDC42 signaling events | 1 |
| NCI-PID | 37006 | PLK1 signaling events | 1 |
| NCI-PID | 37011 | Lissencephaly gene (LIS1) in neuronal migration and development | 2 |
| NCI-PID | 37013 | Ephrin B reverse signaling | 2 |
| NCI-PID | 37014 | Syndecan-2-mediated signaling events | 2 |
| NCI-PID | 37015 | Neurotrophic factor-mediated Trk receptor signaling | 1 |
| NCI-PID | 37017 | Calcineurin-regulated NFAT-dependent transcription in lymphocytes | 1 |
| NCI-PID | 37018 | Fc-epsilon receptor I signaling in mast cells | 2 |
| NCI-PID | 37020 | VEGFR1 specific signals | 1 |
| NCI-PID | 37023 | IL2-mediated signaling events | 1 |
| NCI-PID | 37025 | Syndecan-4-mediated signaling events | 1 |
| NCI-PID | 37027 | BCR signaling pathway | 2 |
| NCI-PID | 37033 | Integrins in angiogenesis | 1 |
| NCI-PID | 37034 | EPO signaling pathway | 2 |
| NCI-PID | 37035 | BMP receptor signaling | 2 |
| NCI-PID | 37038 | EPHB forward signaling | 2 |
| NCI-PID | 37040 | EPHA forward signaling | 2 |
| NCI-PID | 37041 | IL6-mediated signaling events | 1 |
| NCI-PID | 37043 | HIV-1 Nef: Negative effector of Fas and TNF-alpha | 1 |
| NCI-PID | 37044 | TCR signaling in naive CD4+ T cells | 3 |
| NCI-PID | 37046 | Syndecan-3-mediated signaling events | 2 |
| NCI-PID | 37048 | Insulin-mediated glucose transport | 1 |
| NCI-PID | 37050 | Hedgehog signaling events mediated by Gli proteins | 1 |
| NCI-PID | 37051 | ErbB4 signaling events | 1 |
| NCI-PID | 37054 | RhoA signaling pathway | 1 |
| NCI-PID | 37056 | LPA4-mediated signaling events | 1 |
| NCI-PID | 36885 | Signaling events mediated by Hepatocyte Growth Factor Receptor (c-Met) | 2 |
| NCI-PID | 36948 | Ephrin A reverse signaling | 2 |
| NCI-PID | 36972 | Alpha-synuclein signaling | 2 |
| NCI-PID | 37045 | FGF signaling pathway | 1 |

**Supplementary Table -.**
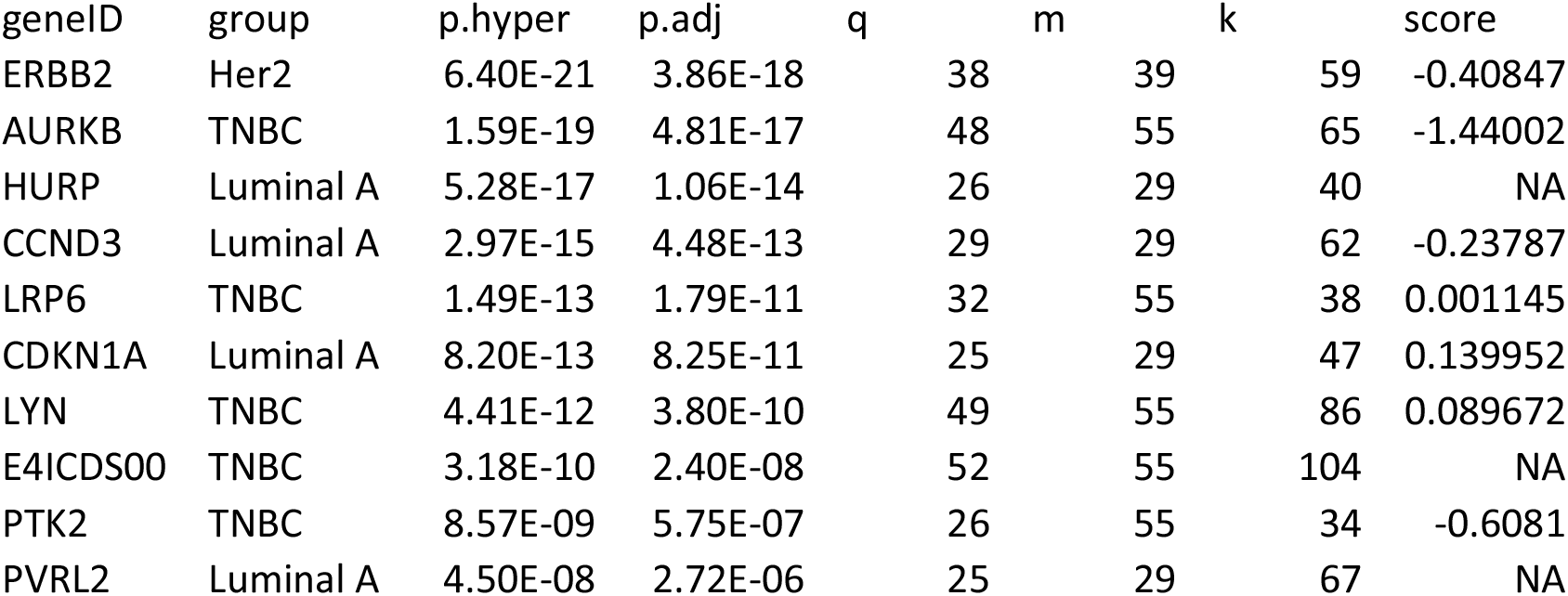
BRCA essentiality results (subtypes). q: number of double positives, m: number of group patients, k: number of patients predicted as essential. Total number of samples = 164. Score: average Achilles score across all cell lines.

| geneID | group | p.hyper | p.adj | q | m | k | score |
| --- | --- | --- | --- | --- | --- | --- | --- |
| ERBB2 | Her2 | 6.40E-21 | 3.86E-18 | 38 | 39 | 59 | -0.40847 |
| AURKB | TNBC | 1.59E-19 | 4.81E-17 | 48 | 55 | 65 | -1.44002 |
| HURP | Luminal A | 5.28E-17 | 1.06E-14 | 26 | 29 | 40 | NA |
| CCND3 | Luminal A | 2.97E-15 | 4.48E-13 | 29 | 29 | 62 | -0.23787 |
| LRP6 | TNBC | 1.49E-13 | 1.79E-11 | 32 | 55 | 38 | 0.001145 |
| CDKN1A | Luminal A | 8.20E-13 | 8.25E-11 | 25 | 29 | 47 | 0.139952 |
| LYN | TNBC | 4.41E-12 | 3.80E-10 | 49 | 55 | 86 | 0.089672 |
| E4ICDS00 | TNBC | 3.18E-10 | 2.40E-08 | 52 | 55 | 104 | NA |
| PTK2 | TNBC | 8.57E-09 | 5.75E-07 | 26 | 55 | 34 | -0.6081 |
| PVRL2 | Luminal A | 4.50E-08 | 2.72E-06 | 25 | 29 | 67 | NA |

